# Fibril elongation by human islet amyloid polypeptide is the main event linking aggregation to membrane damage

**DOI:** 10.1101/2022.10.14.512241

**Authors:** Barend O.W. Elenbaas, Stefanie M. Kremsreiter, Lucie Khemtemourian, J. Antoinette Killian, Tessa Sinnige

**Affiliations:** Membrane Biochemistry and Biophysics, Bijvoet Centre for Biomolecular Research, Utrecht University, Padualaan 8, 3584 CH, Utrecht, The Netherlands; Institute of Chemistry & Biology of Membranes & Nanoobjects (CBMN), CNRS UMR5248, University of Bordeaux, Bordeaux INP, allée Geoffroy St-Hilaire, 33600, Pessac, France

## Abstract

The aggregation of human islet amyloid polypeptide (hIAPP) is linked to the death of pancreatic β-cells in type II diabetes. The process of fibril formation by hIAPP is thought to cause membrane damage, but the precise mechanisms are still unclear. Previously, we showed that the aggregation of hIAPP in the presence of membranes containing anionic lipids is dominated by secondary nucleation events, which occur at the interface between existing fibrils and the membrane surface. Here, we used vesicles with different lipid composition to explore the connection between hIAPP aggregation and vesicle leakage. We found that different anionic lipids promote hIAPP aggregation to the same extent, whereas remarkably stochastic behaviour is observed on purely zwitterionic membranes. Vesicle leakage induced by hIAPP consists of two distinct phases for any of the used membrane compositions: (i) an initial phase in which hIAPP binding causes a certain level of leakage that is strongly dependent on osmotic conditions, membrane composition and the used dye, and (ii) a main leakage event that we attribute to elongation of hIAPP fibrils, based on seeded experiments. Altogether, our results shed more light on the relationship between hIAPP fibril formation and membrane damage, and strongly suggest that oligomeric intermediates do not considerably contribute to vesicle leakage.

## Introduction

Type II diabetes poses a growing health problem to society, affecting increasing numbers of patients^1,2^. The disease is characterized by the death of the insulin-producing β-cells in the pancreatic islets^3^. The death of these cells is strongly associated with the formation of human islet amyloid polypeptide (hIAPP) aggregates^4,5^. hIAPP is a 37 amino acid long peptide, which the β-cells co-produce and secrete together with insulin in response to glucose. Characteristic of amyloids, hIAPP forms fibrils with a cross-β structure^6–8^. The process of fibril formation is thought to disturb cellular membranes, causing toxicity and ultimately cell death, but the precise mechanisms remain unclear.

Various hypotheses have been proposed on the relationship between hIAPP aggregation and membrane damage. Fibril growth has been suggested to induce vesicle leakage by membrane deformation^9,10^, likely accompanied by the extraction of lipids^11^. Biphasic leakage profiles have also been reported, in which the first phase was proposed to stem from small oligomers and the second phase from fibril growth^12^. The relevance of leakage assays as a measure for cellular toxicity was questioned after IAPP from rat and mice (here abbreviated as mIAPP) was also found to induce rapid leakage of model membranes^13^. This was unexpected because mIAPP, which differs from hIAPP at six positions, is non-amyloidogenic^14^ and non-toxic^15^. In a study using cell-derived giant unilamellar vesicles (GUVs), also both hIAPP and mIAPP induced a first phase of leakage^16^. hIAPP in addition caused a second phase of leakage, which was in this case postulated to arise from non-amyloid, large oligomeric species.

Others have investigated the mechanism of membrane disruption by looking directly at cellular toxicity. The addition of soluble hIAPP, but not of mature aggregates, was found to be toxic to islet cells^17^. When hIAPP was allowed to aggregate *in vitro* and added to cultured cells at different timepoints, early fractions containing oligomers were found to be most toxic^18^. However, mIAPP also has the propensity to oligomerise in solution^18^ and on membranes^19^, despite the fact that it is not toxic. Altogether, no unifying model has emerged to explain the relationship between hIAPP aggregation and toxicity.

Detailed knowledge of the aggregation mechanism of hIAPP in the presence of membranes is essential to make further progress. hIAPP aggregation is accelerated by membranes containing anionic lipids^19–23^. Recently, we established a molecular mechanism by which the anionic lipid 1,2-dioleoyl-*sn*-glycero-3-phospho-L-serine (DOPS) catalyses hIAPP fibril formation^23^. We showed that this lipid strongly promotes both primary and secondary nucleation events, but not fibril elongation. Nevertheless, elongation must occur at least in part on the membrane surface, because secondary nucleation takes place at the interface of the lipid head groups and existing fibrils. The question which of these processes is responsible for membrane damage thus remained unanswered.

Here, we compare the aggregation kinetics and vesicle leakage for different membrane compositions, with the goal of identifying the molecular event that is most closely associated with membrane damage in a physiological context.

## Results

### hIAPP aggregation kinetics depend only on the charge of the lipid head group

Previously, we reported that the anionic lipid PS promotes nucleation of hIAPP on the membrane^23^. We first set out to determine whether this effect depends on the chemical properties of the lipid head group, or whether it is simply due to electrostatic interactions with the positively charged hIAPP peptide. We performed Thioflavin T (ThT) aggregation assays of synthetic hIAPP with amidated C-terminus in the presence of large unilamellar vesicles (LUVs) of different lipid compositions at a constant lipid concentration (**Fig. 1**, **Fig. S1**). Lipids were present in excess to avoid the aggregation of hIAPP in solution, which would change the overall aggregation kinetics^23^.

**Figure 1.**
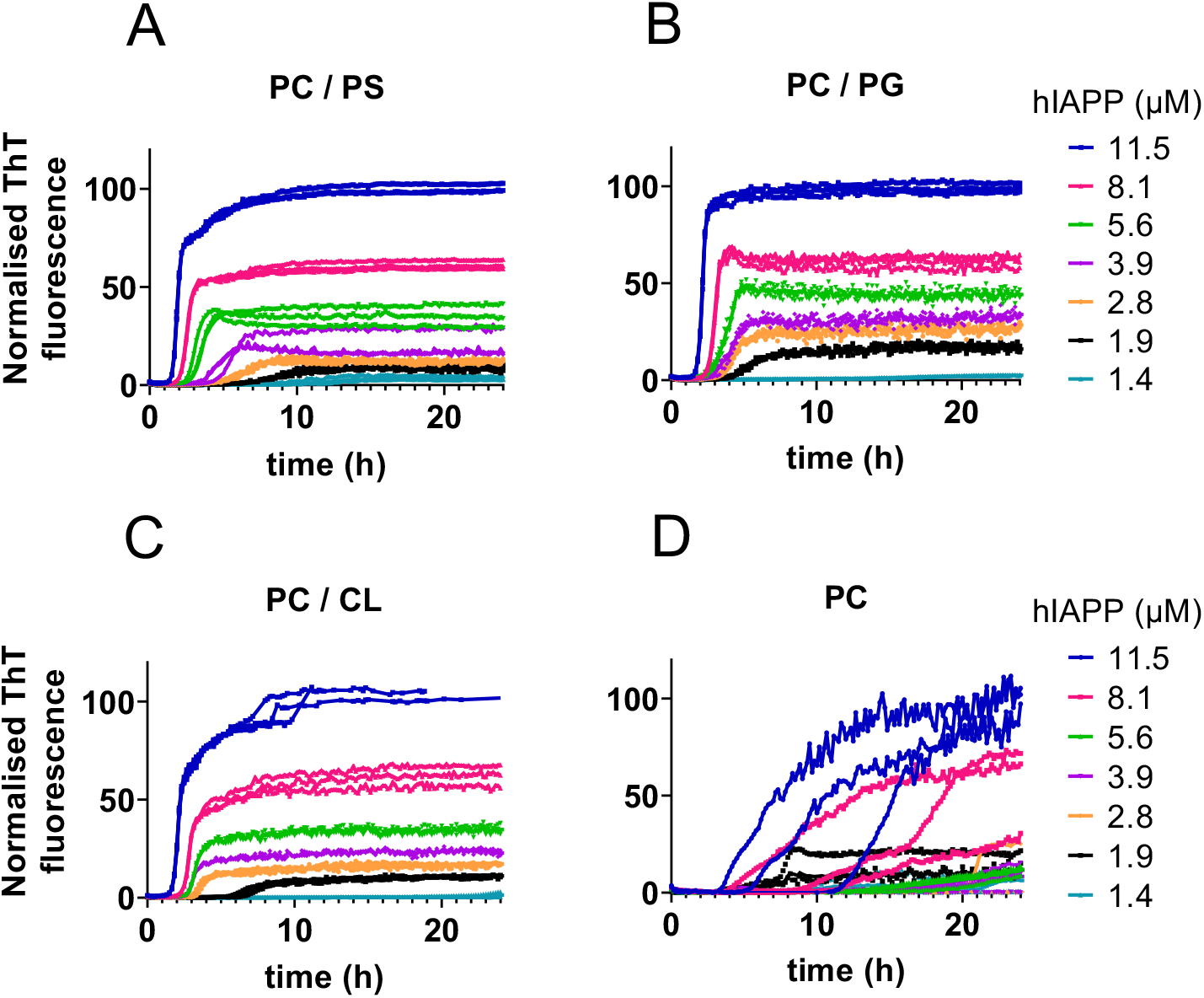
hIAPP aggregation kinetics in the presence of LUVs with four distinct lipid compositions as followed by ThT fluorescence. (A) PC/PS, (B) PC/PG, (C) PC/CL and (D) PC. The concentration of lipids was kept constant at 863 μM on phosphate basis. The signals are normalised to the maximal fluorescence of the highest (11.5 μM) hIAPP concentration. Each hIAPP concentration was measured in triplicate, shown as individual traces.

We first inspected hIAPP aggregation in the presence of vesicles containing different anionic lipids (**Fig. 1A, B, C**). Phosphatidylglycerol (PG) has been used previously in hIAPP aggregation studies^19^. PG has different properties from PS, having a smaller head group with a different number of total charges. Nevertheless, it promotes hIAPP aggregation to a similar extent as PS in our assay (**Fig. 1A, B**). To test the effect of local charge distribution, we also examined the phospholipid cardiolipin (CL). This lipid carries a total of four acyl chains and two negative charges per molecule, and has an even smaller relative head group size than PG. CL was introduced at half the molar ratio to achieve the same net negative surface charge on the membrane as for the single anionic head groups, again resulting in kinetic profiles that look highly similar (**Fig. 1C**). Thus, the aggregation propensity of hIAPP is independent of the charge distribution or the specific lipid head group chemistry.

### hIAPP aggregates stochastically on zwitterionic membranes

The kinetics of hIAPP aggregation in the presence of purely zwitterionic phosphatidylcholine (PC) lipids are very different from those in the presence of anionic lipids (**Fig. 1D**). We encountered high variability between replicates with most datasets showing an extended lag phase, but occasionally we also observed very rapid aggregation (**Fig. S2A, B**). The variable results for PC were persistent, even though the experiments were performed in parallel with the other lipid compositions. We therefore believe that hIAPP intrinsically displays stochastic aggregation behaviour under these conditions. This could be the result of a very low nucleation propensity, related to the weak and transient binding to these zwitterionic membranes^19,24^. At the same time, the traces look different from those observed for the same hIAPP preparation in the absence of LUVs (**Fig. S2C**), indicating that hIAPP nucleation takes place primarily on the PC membrane rather than in solution. The sigmoidal profile of hIAPP aggregation in the presence of PC demonstrates that the auto-catalytic process of secondary nucleation still dominates under these conditions, similarly as for anionic membranes^23^.

### Conformation of hIAPP is similar in the presence of membranes with different anionic lipids

To further explore the difference between hIAPP binding to purely zwitterionic or anionic membranes, we turned to circular dichroism (CD) spectroscopy. hIAPP has been reported to adopt a partial a-helical structure upon binding to membranes containing PS or PG^19,21,25,26^. Using the same lipid compositions as in the ThT assays, we observed that the CD spectra of hIAPP initially display an a-helical signature in the presence of membranes containing any of the anionic lipids PS, PG or CL (**Fig. 2A, B, C and F**). By contrast, the spectrum of hIAPP in the presence of pure PC vesicles has the same random coil appearance as hIAPP in solution (**Fig. 2D, E and F**), suggesting that hIAPP binding to these membranes is weak or transient. Over time, all conditions lead to a clear shift towards β-sheet content (**Fig. 2A-E**), indicative of fibril formation in agreement with previous studies^19,21,25,26^.

**Figure 2.**
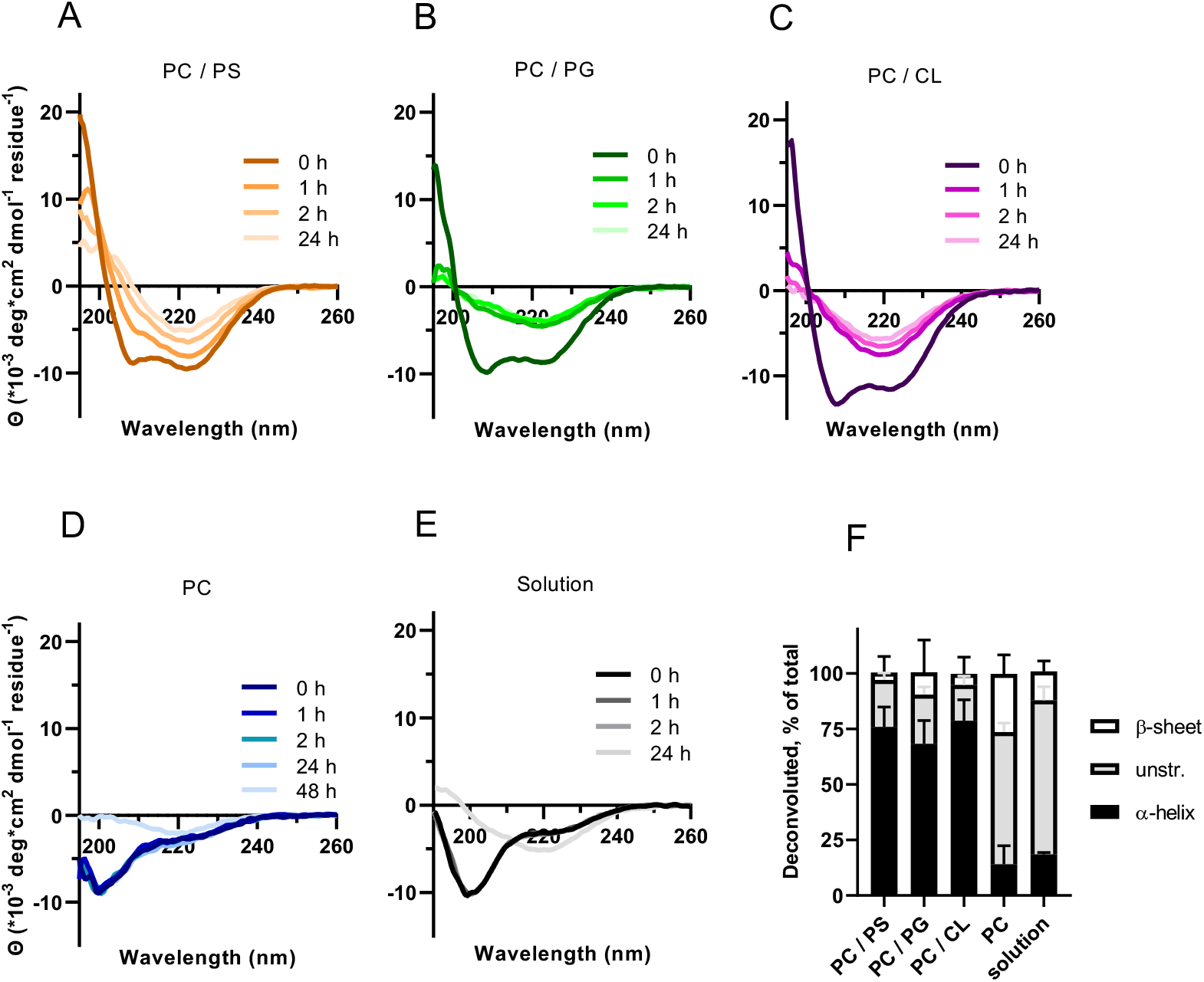
CD spectra of 25 μM of hIAPP in the presence of various lipid composition at 1:50 molar peptide:lipid ratio, and in solution. (A) PC/PS, (B) PC/PG, (C) PC/CL, (D) 100% PC and (E) without lipids. Over time, the spectra shift towards β-sheet conformation (minimum at 218 nm). (F) Deconvolution of the spectra at t = 0, showing predominantly a-helical conformation for PC/PS, PC/PG, and PC/CL, and unstructured for PC and solution.

CD spectra of mIAPP display similar characteristics as those of hIAPP, but remain stable over time given the lack of amyloid formation by this variant (**Fig. S3**). Altogether these data indicate that the initial interaction of hIAPP and mIAPP with membranes containing anionic lipids is comparable, whether the lipid is PS, PG or CL. For hIAPP, this is consistent with the aggregation behaviour across the different anionic lipids tested (**Fig. 1A, B and C**). For mIAPP, these data provide confidence that this variant can be used to discriminate binding from aggregation, as detailed in the following.

### hIAPP induced vesicle leakage shows two-phase behaviour

Next, we used calcein leakage assays to explore the correlation between hIAPP fibril formation and membrane damage (**Fig. 3**). ThT assays were run in parallel to minimize the variation in experimental conditions. The leakage assays in the presence of hIAPP show a striking two-phase behaviour, consistent with previous reports^9,12,13^ (**Fig. 3A-D**). The first phase starts directly at time zero, when no ThT signal is visible yet (**Fig. 3E-H**). The second phase is responsible for the largest fraction of leakage, and corresponds to the growth phase of the ThT curve during which the fibril mass accumulates (**Fig. 3I**).

**Figure 3.**
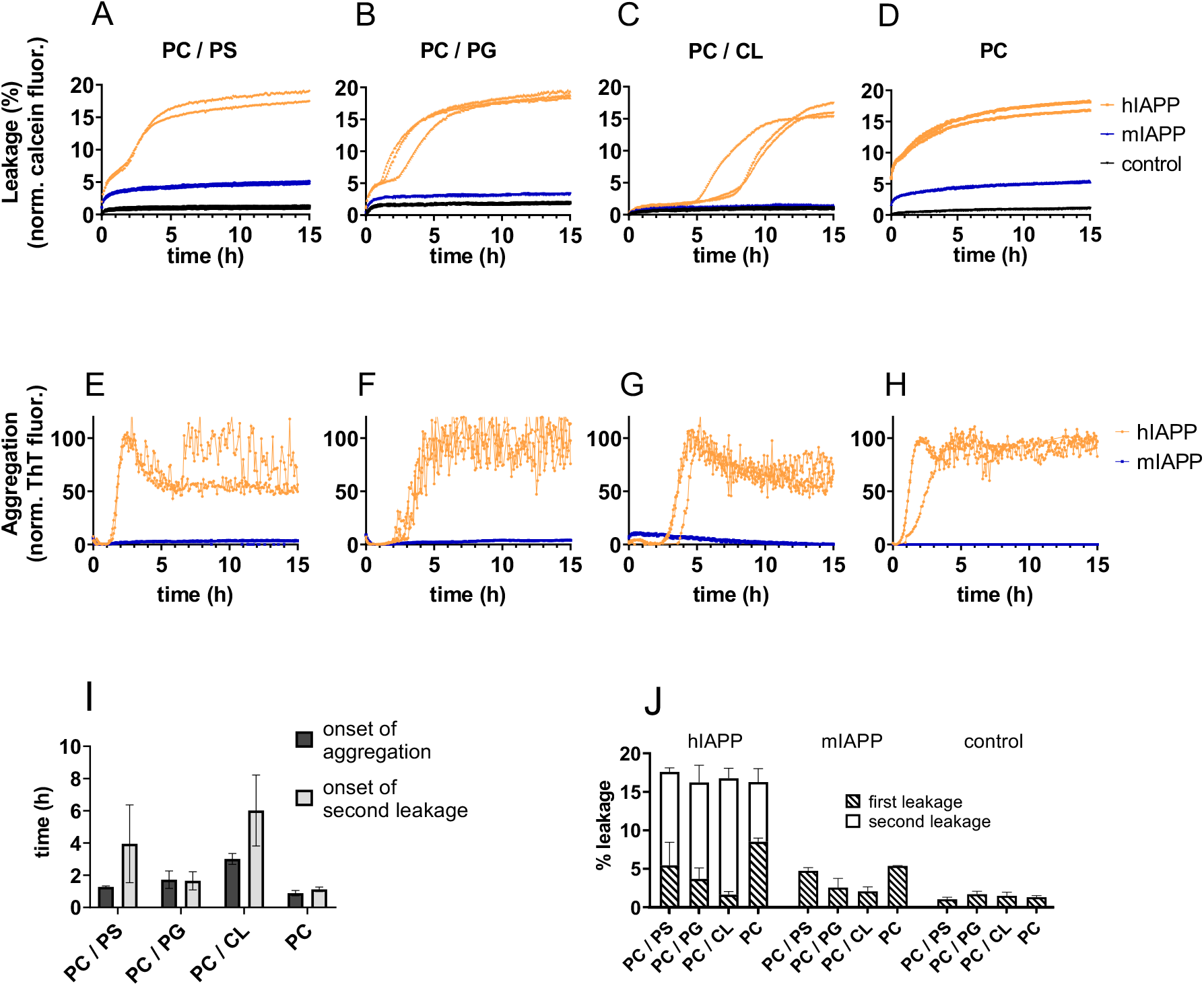
Parallel assay measuring calcein leakage (A-D) and ThT aggregation (E-H) for four different membrane compositions: (A, E) PC/PS, (B, F) PC/PG, (C, G) PC/CL, and (D, H) PC. The fluorescence in the presence of hIAPP is shown in orange, mIAPP in blue and milliQ in black. Individual traces of triplicate measurements are shown. (I) The onset of hIAPP aggregation (black) and that of the second phase of leakage (grey). Onset of aggregation was defined as the time point at which the fluorescence is 5% above the baseline; the inflection points in the calcein curves were taken as the start of the second leakage event. (J) The percentage leakage for hIAPP, mIAPP and milliQ (control), with respect to full solubilization by Triton X-100. The data in (I) and (J) are compiled from two independent experiments each performed in triplicate.

The non-aggregating mIAPP also shows an initial leakage phase starting at time zero for most of the lipid compositions tested (**Fig. 3A-D**). Although the percentage of leakage is somewhat lower than for hIAPP (**Fig. 3J**), this result suggests that the first phase of membrane damage is caused by binding of non-toxic peptide species. Vesicles containing CL appear more resistant against this type of damage (**Fig. 3C, J**). This difference is most likely due to an increased stability of CL containing membranes, since the mode of interaction of hIAPP is similar for all anionic lipids as inferred from the ThT and CD experiments.

Even though the interaction of hIAPP with pure PC membranes is weaker and no stable α-helix formation is observed in the CD experiments (**Fig. 2D**), the zwitterionic vesicles are affected by leakage to a similar extent as those containing anionic lipids (**Fig. 3D, J**). The early leakage event is even somewhat more pronounced, suggesting that transient interactions with hIAPP as well as mIAPP are sufficient to induce this type of membrane permeability. Again, we observed high variability between experiments for pure PC vesicles. The dataset shown in **Fig. 3D** and **H** hardly has a lag phase, whereas other independent experiments display extended lag times before the second phase of leakage (**Fig. S4**).

### Initial leakage event depends on experimental conditions

The initial phase of leakage has previously been observed by several labs, although to different extents^9,12,13^. The fact that also mIAPP induces this phenomenon has led to the suggestion that toxicity and membrane leakage are not connected^13^. Inspired by our observation that the initial leakage is less pronounced for the more stable CL membranes, we probed to what extent it can be tuned by experimental conditions (**Fig. 4**). First, we tested the effect of osmotic pressure. We observed a strong correlation between leakage and differences in osmolality between the outside and inside of the vesicles, with higher osmotic pressure inside the vesicles leading to increased leakage (**Fig. 4A, C**). Using dynamic light scattering (DLS) we found that the size of the vesicles is only slightly affected by the osmotic differences (**Table S1**). However, the osmotic differences are clearly sufficient to destabilize the vesicles and cause differences in initial leakage.

**Figure 4.**
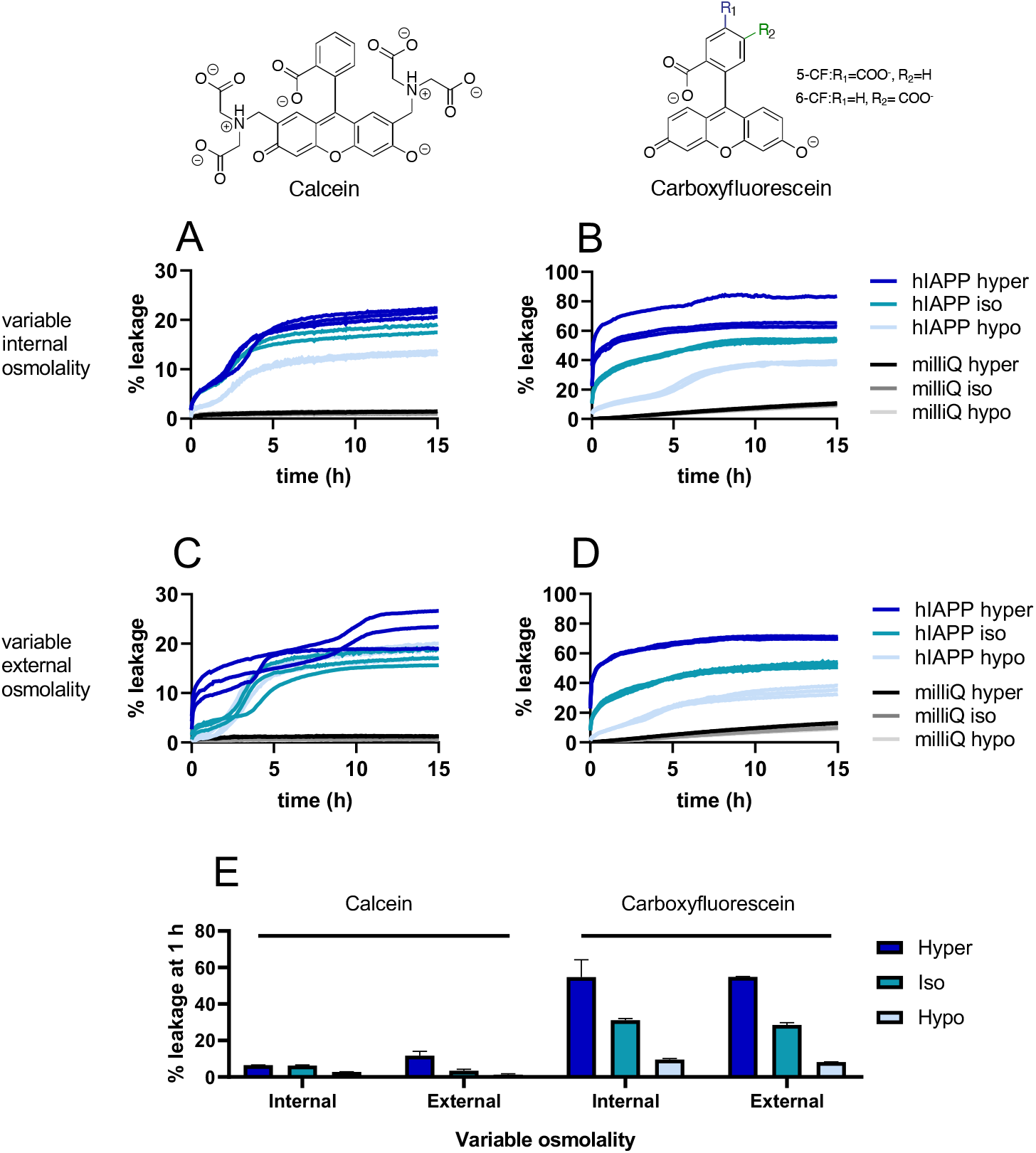
Leakage of PC/PS vesicles induced by hIAPP under various osmotic conditions. (A, C) Calcein fluorescence. (B, D) Carboxyfluorescein fluorescence. Either the internal osmolality (A, B) or the external osmolality (C, D) is varied. Hyperosmotic (dark blue), isoosmotic (turquoise) and hypoosmotic (light blue) refer to the conditions inside the vesicles. Controls were performed by adding milliQ water (black and grey). Lines show the separate traces of triplicate measurements. (E) Quantification of the leakage after 1 h incubation for the different conditions.

Moreover, we found that vesicles filled with carboxyfluorescein show markedly higher levels of leakage than those loaded with calcein, the latter being a bulkier molecule (**Fig. 4B, D, E**). Notably, in the leakage experiments using carboxyfluorescein and iso- or hyperosmotic conditions, the biphasic behaviour is no longer visible due to the overwhelming magnitude of the initial leakage event (**Fig. 4B, D**). On the other hand, initial leakage is completely abolished under hypoosmotic conditions in combination with the larger calcein dye (**Fig. 4C**). Together with our earlier result that mIAPP similarly causes the first phase of leakage, we conclude that this phenomenon is the result of transient membrane permeabilization caused by IAPP binding. However, only the second phase of leakage is associated with the aggregation pathway of hIAPP.

### Second leakage phase is caused by fibril elongation

The second phase of leakage corresponds to the growth phase of the ThT curve (**Fig. 3**), which is dominated by secondary nucleation and elongation. To dissect which of these two processes is responsible for the bulk of the calcein leakage observed, we performed experiments in the presence of high concentrations of seeds (**Fig. 5**). Adding preformed fibril seeds bypasses the need for primary nucleation, and at very high seed concentrations the abundance of fibril ends causes elongation to dominate over secondary nucleation at the immediate start of the experiment^27^. These conditions are therefore also expected to eliminate the potential formation of on-pathway oligomers.

**Figure 5.**
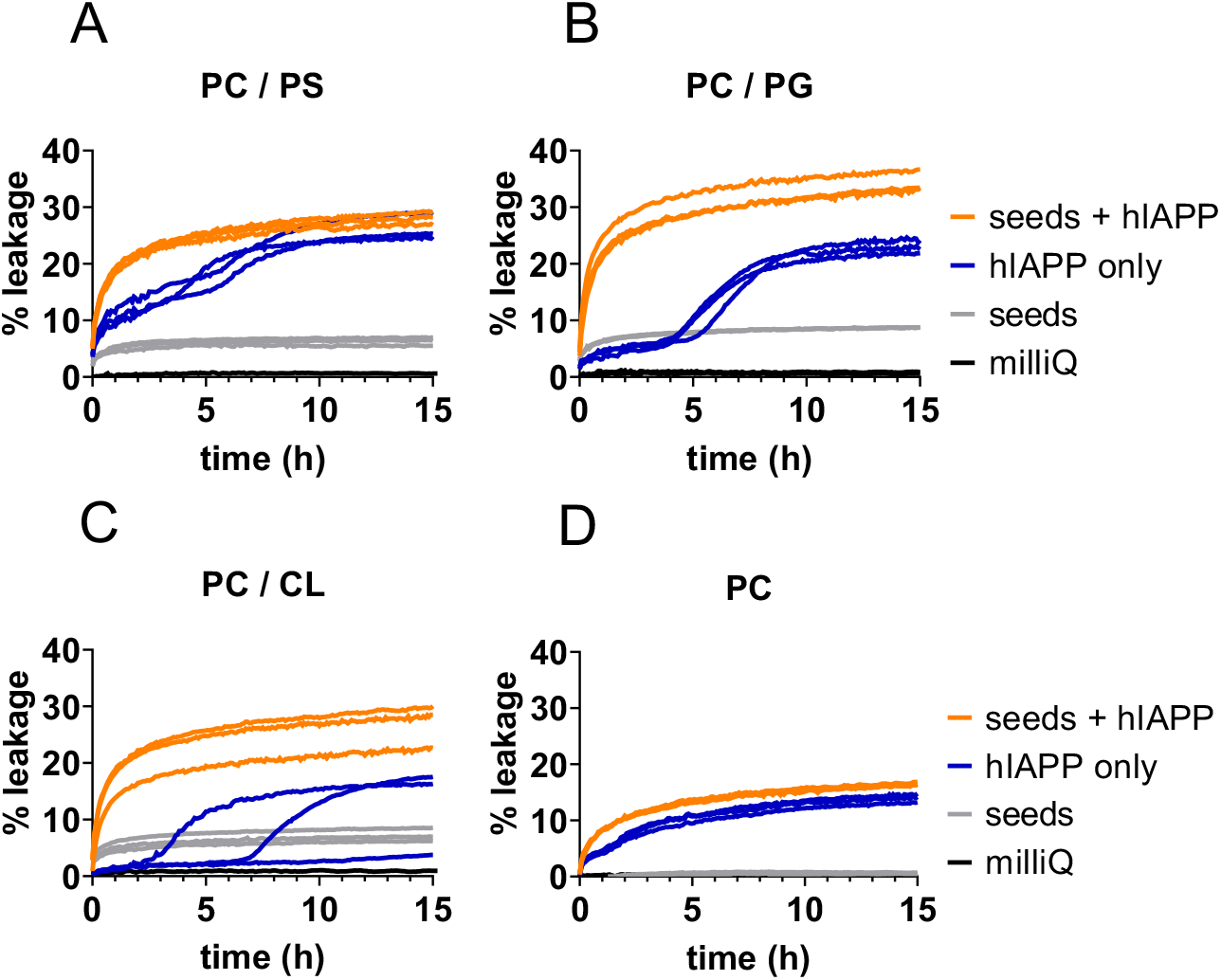
Seeded leakage assays of calcein vesicles with four different lipid compositions: (A) PC/PS, (B) PC/PG, (C) PC/CL and (D) PC vesicles. The unseeded leakage was induced by 5 μM hIAPP (blue). The seeded leakage was induced by 5 μM of hIAPP plus 2.14 μM seeds (monomer based), which amounts to a 30 % seeded reaction (orange). 2.14 μM seeds (grey) and milliQ only (black) were used as controls. Individual traces of triplicate measurements are shown.

In the presence of monomeric hIAPP with 30% fibril seeds added, we found that leakage progresses directly from time zero, without the two-phase behaviour observed in the absence of seeds (**Fig. 5**). The seeds alone induce low levels of leakage, which may be due to their interaction with the membrane. These data altogether suggest that hIAPP fibril elongation is the main process that disturbs membrane integrity.

## Discussion

How hIAPP aggregation relates to toxicity and β-cell death is the subject of a long-standing debate. Here, we demonstrate that fibril elongation of hIAPP is the main aggregation step responsible for the leakage of synthetic membranes with various lipid compositions.

Previously, we reported that the anionic lipid PS strongly accelerates both primary and secondary nucleation of hIAPP in PC/PS model membranes^23^. Any of the negatively charged lipids we tested in this work has a similar effect on the aggregation kinetics. Thus, the catalytic effect does not appear to be mediated by the specific chemistry of the lipid head group, but may simply be a result of increased binding affinity due to electrostatic interactions. The dissociation constant of hIAPP binding to pure PC LUVs was estimated to be 1.4 mM, whereas incorporation of 20% PS lowers the Kd to 200 μM^24^. Stronger binding increases the fraction of hIAPP that is bound to the membrane, thus increasing the probability of nucleation.

We find that hIAPP also aggregates on purely zwitterionic membranes, but with remarkably stochastic behaviour. This is unlikely to be an experimental artefact, given the high reproducibility of the data acquired in the presence of anionic lipids, which we report here and in earlier work^23^. The rate constant for primary nucleation of hIAPP on PC/PS membranes was found to be orders of magnitude lower than that for secondary nucleation^23^. We speculate that primary nucleation of hIAPP on PC membranes is even more infrequent, due to the transient nature of the interaction. However, once the first nucleation events have happened, fibril formation will proceed readily due to the auto-catalytic nature of the process.

Our vesicle leakage data show that i) hIAPP induced leakage is biphasic, and ii) both hIAPP and mIAPP induce the initial phase of leakage. Previous reports of these phenomena led to conflicting interpretations^12,13,16^, which we here resolve. We suggest that the first leakage phase, preceding detectable fibril formation, stems from interactions of both hIAPP and mIAPP with the membrane that cause transient perturbations. Even PC, which has much lower affinity for IAPP than the anionic membranes, shows pronounced initial leakage, suggesting that weak interactions are sufficient to induce this type of permeation.

The initial leakage event strongly depends on experimental conditions, including osmotic pressure and the used dye. Based on our results we recommend to verify osmolality of the vesicles and the outside buffer, and to use the larger dye calcein, rather than carboxyfluorescein, when investigating membrane damage in relation to protein aggregation. Cellular membranes are likely to be much more rigid than the model vesicles used here, as they are full of proteins. Interestingly, we observed that vesicles containing CL, which confers increased stability as a double lipid, do not display the initial phase of calcein leakage under isoosmotic conditions. Taken together, our data suggest that the initial leakage event may not be physiologically relevant.

The second phase of leakage is more likely to be related to the mechanism of toxicity, as it is only observed for hIAPP, but not for mIAPP, and corresponds to the growth phase of the ThT curve. Using seeded experiments, we demonstrate that fibril elongation is responsible for this event. Notably, elongation does not depend on the presence of anionic lipids, as we have shown earlier^23^. Consistent with these results, zwitterionic PC membranes display similar levels of fibrillisation-induced leakage as the anionic membranes. Thus, we suggest that membranes of any composition can succumb to damage by hIAPP fibril elongation, once initial nucleation events have taken place.

Our data argue against a role for on-pathway oligomers in membrane permeation. The nucleation events that would give rise to such species are bypassed in the highly seeded leakage experiments. We would like to point out that the addition of soluble hIAPP to cultured cells is very likely to lead to fibril growth on the plasma membrane, offering an alternative explanation for the toxicity previously attributed to early oligomers.

## Materials and Methods

### Materials

Human islet amyloid polypeptide (hIAPP) (Amylin (human) trifluoroacetate salt, Lot no. 1000026803) and murine IAPP (Amylin (mouse, rat) trifluoroacetate salt, Lot no. 1000038853) were purchased from Bachem. The peptides have a disulfide bond and are amidated at the C-terminus. All phospholipids were purchased from Avanti Polar Lipids: 1,2-dioleoyl-*sn*-glycero-3-phosphocholine (DOPC), 1,2-dioleoyl-*sn*-glycero-3-phospho-L-serine (DOPS), 1,2-dioleoyl-*sn*-glycero-3-phospho-(?-rac-glycerol) (sodium salt) (DOPG) and 1’,3’-bis[1,2-dioleoyl-*sn*-glycero-3-phospho]-glycerol (sodium salt) (cardiolipin, TOCL). Sephadex G50-fine was bought from Pharmacia fine chemicals. The 5(6)-carboxyfluorescein was from Kodak. All other chemicals were from Sigma-Aldrich.

### IAPP preparation

hIAPP and mIAPP were treated identically. The polypeptides were dissolved in hexafluoro-2-propanol to a concentration of 1 mM and the solution was left on the bench overnight at room temperature. The solution was then aliquoted in small (< 50 μL) volumes, which were placed under a high vacuum for 1 h. The aliquots were stored at −80 °C and dissolved prior to use in milliQ water to 160 μM (kinetics experiments), 175 μM (circular dichroism experiments) or 200 μM (leakage experiments).

### hIAPP seed preparation

hIAPP seeds were prepared by taking an hIAPP aliquot prepared as described above from the −80 °C freezer. The peptide was dissolved to a concentration of 50 μM in 10 mM Tris-HCl pH 7.4, 100 mM NaCl. The solution was left on the bench for 24 h to allow for fibril formation, and subsequently sonicated four times for one minute in a Branson 3800 bath sonicator. Seeds were used directly after preparation.

### Vesicle preparation

Stock solutions of the lipids were made in chloroform and the phosphate concentration was determined using a Rouser assay^28^. The molar ratio was 7:3 for DOPC:DOPS and DOPC:DOPG, and 14:3 for DOPC:TOCL. The appropriate volumes of each lipid solution were mixed and dried under a nitrogen gas flow in a 42 °C water bath to evaporate the solvent. The samples were placed in a desiccator for at least 30 min to remove any residual solvent. The lipid film was then resuspended in 10 mM Tris-HCl pH 7.4, 100 mM NaCl to allow for spontaneous formation of multilamellar vesicles (MLVs). The suspensions were left on the bench at room temperature for 1 h with gentle swirling every 10 min to suspend the lipid film. The MLVs were homogenized with 10 freeze thaw cycles by alternatingly placing them in ethanol cooled by dry ice pellets, and lukewarm water. To create large unilamellar vesicles (LUVs), MLVs were extruded through a 200 nm pore filter (Anotop 10, Whatman), using an Avanti Polar Lipids extruder set. This was done 10 times back and forth, followed by one more passage to ensure that the vesicles ended on the opposite side from where they initiated. The final lipid concentration of the LUV suspension was determined again using a Rouser assay^28^. LUVs were stored at 4 °C and used within one week of preparation.

### Thioflavin T assay

Thioflavin T (ThT) assays were performed as described previously^23^. All assays were performed at a constant lipid concentration of 863 μM on phosphate basis, leading to a minimal peptide:lipid ratio of 1:75 for the highest hIAPP concentration of 11.5 μM.

### CD spectroscopy

CD spectra of hIAPP and mIAPP were recorded on a Jasco 810 spectropolarimeter. The temperature was set at 25 °C with a Jasco CDF-426S element. Quartz cells (Hellma GmbH) were used with a path length of 1 mm and an internal volume of 350 μL. The peptides were dissolved in milliQ and then diluted to a final concentration of 25 μM in 10 mM sodium phosphate buffer pH 7.4 with or without LUVs added. LUVs were prepared as described above, but in 10 mM sodium phosphate pH 7.4. The final peptide:lipid ratio was 1:50.

The first measurement was started immediately after the IAPP was added to the buffer, with subsequent measurements at regular time intervals. The CD spectra were measured in 0.2 nm intervals over a wavelength range of 190 – 270 nm with a scanning speed of 50 nm/min. Each spectrum was the average of five scans. A baseline correction was carried out by subtracting the average signal between 260-270 nm from the entire spectrum. Deconvolution of the secondary structure content was performed using CDPro. The values from three algorithms (SELCON3, CDSSTR and CONTIN) were averaged.

### Standard leakage assay

The preparation of the vesicles was identical for calcein and carboxyfluorescein. Lipid films were created as described above, but resuspended in fluorophore solution with an osmolality of 250 mOsm/kg. Stock solutions were prepared by dissolving 100 mM dye in milliQ water using a few drops of 10 M NaOH. The pH was brought back to 7.4 using 1 M HCl. The stock was filtered through a 200 nm cellulose acetate membrane and the osmolality was measured using a K-7400S Semi-Micro Osmometer (KNAUER). The solution was diluted until a value of 250 (± < 5 %) mOsm/kg was reached.

The vesicles were homogenised and made into LUVs as described above. To remove the dye outside of the LUVs, a 10 mL gravity size exclusion column was used. The Sephadex resin was soaked in outside buffer, which was set to an osmolality of 256 mOsm/kg (10 mM Tris-HCl pH 7.4, 118 mM NaCl) to compensate for the slight dilution by the addition of hIAPP or control solutions. After elution from the column, the phosphate concentration was measured again using a Rouser assay^28^.

The components were added in a volume ratio of 145:50:5, outside buffer:LUVs:IAPP. Final concentrations were 500 μM lipids and 5 μM IAPP. The positive control with complete leakage was measured by adding 5 μL of 10 % Triton X-100 instead of IAPP. To the negative controls, 5 μL of milliQ water was added. The excitation and emission wavelengths were 475-10 and 515-15 nm with a dichroic mirror setting of 493.8 nm for calcein, and 480-10 and 520-10 nm with a dichroic mirror setting of 500 nm for carboxyfluorescein.

### Leakage assay with variable osmolalities

Vesicles with three different internal osmolalities were used. The procedure was the same as for the standard leakage assay except that the lipid film was suspended in a fluorophore solution with an osmolality of either 300, 250 or 200 mOsm/kg. These were all diluted from the same stock as described above.

To vary external osmolality, vesicles with an internal osmolarity of 250 mOsm/kg were used. Outside buffers with an osmolality of 190, 256 and 328 mOsmol/kg were made by adding either concentrated NaCl or milliQ to the 256 mOsmol/kg buffer solution.

### DLS measurements

A Zetasizer 3000 (Malvern Panalytical) was used for DLS experiments to determine LUV size. The sample concentration was between 5-10 mM of lipids on phosphate basis. Each experiment consisted of three runs with 10 measurements of maximally 120 seconds each and was performed at room temperature. Detection of the backscattering was done at 173°. The data were processed using a mono modal analysis model. The reported hydrodynamic size values were obtained from the number distributions.

## Notes

### Competing Interest Statement

The authors have declared no competing interest.

