## Supplementary information for "Fibril elongation by human islet amyloid polypeptide is the main event linking aggregation to membrane damage"

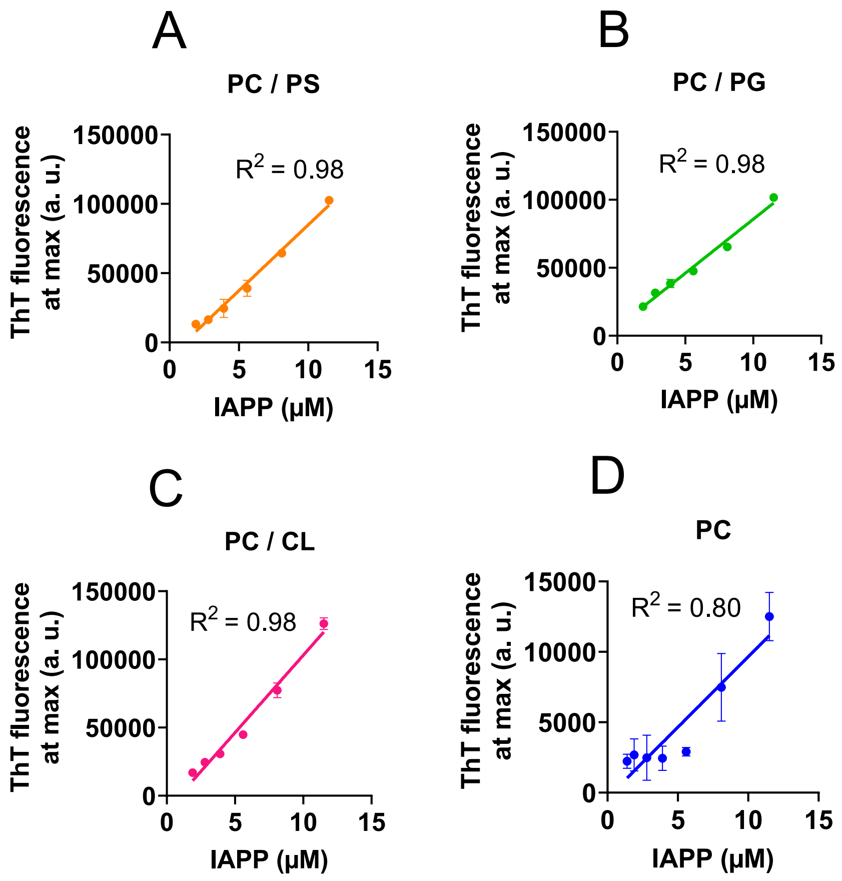


**Figure S1.** Correlation of the maximum fluorescence intensity and the hIAPP concentration for the aggregation kinetics shown in Figure 1. (A) PC/PS, (B) PC/PG, (C) PC/CL and (D) PC. Note that the absolute ThT signal is about 10-fold higher for the anionic membranes than for PC only, which may be related to the positive charge of ThT. The reduced correlation between the hIAPP concentration and the ThT signal in the presence of PC membranes (D) can be attributed to the fact that the plateau was not yet reached for some of the datapoints.

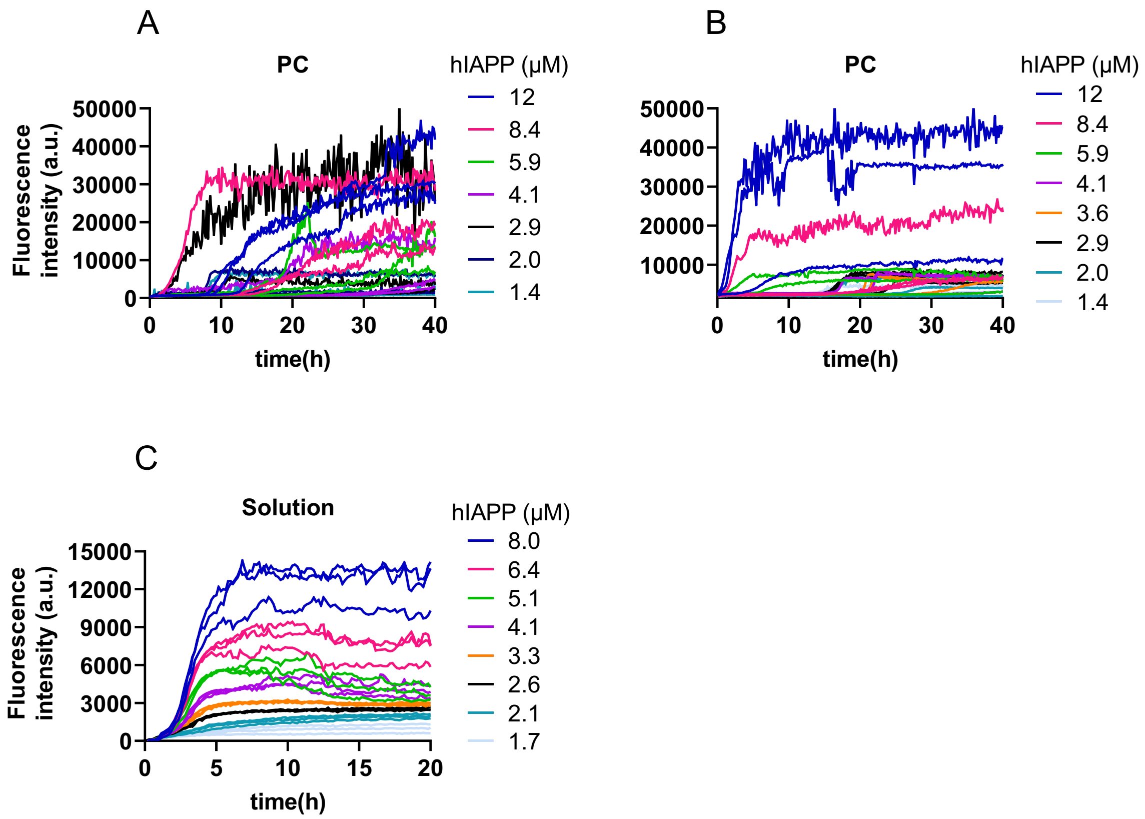


**Figure S2.** Additional ThT aggregation kinetics in the presence of PC and in solution. (A, B) Independent hIAPP aggregation experiments in the presence of PC LUVs. Stochastic aggregation behaviour is apparent even between individual wells in the triplicates. (C) hIAPP aggregation in the absence of LUVs displays very different kinetics, excluding the possibility that hIAPP does not bind to PC membranes leading to its aggregation in solution.

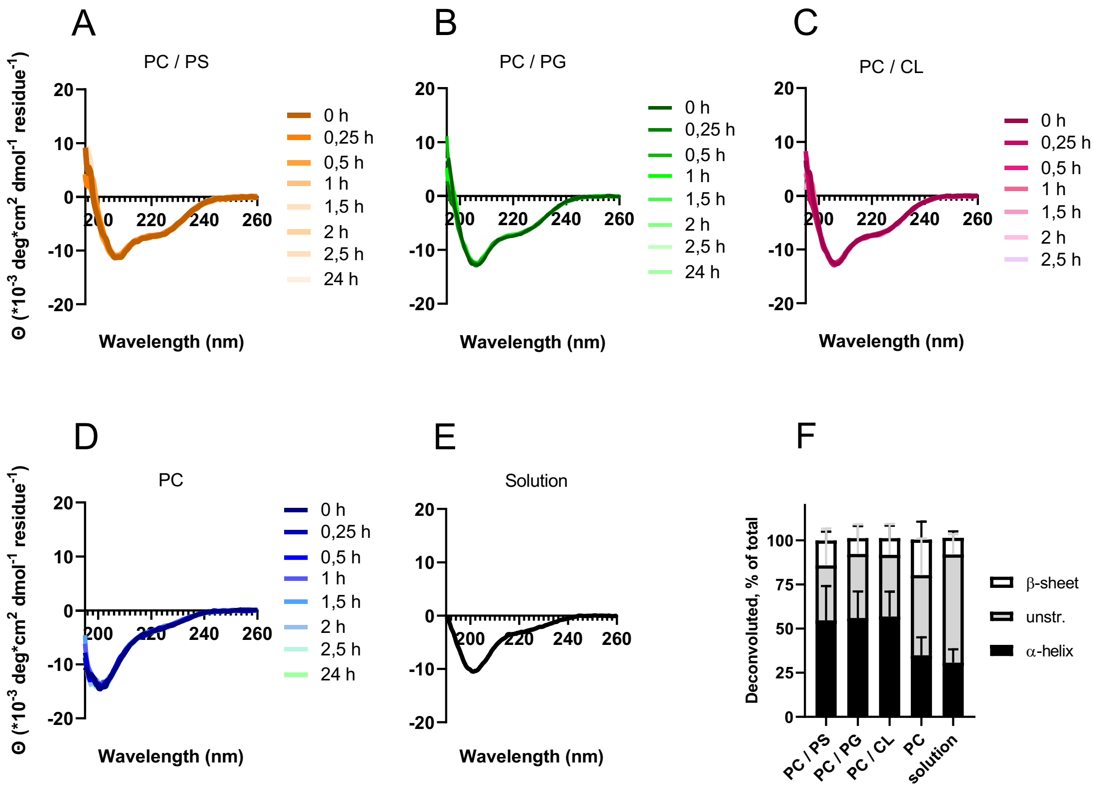


**Figure S3.** Circular dichroism spectra of 25 µM mIAPP in the presence of various lipid composition at 1:50 molar peptide:lipid ratio, and in solution. (A) PC/PS, (B) PC/PG, (C) PC/CL, (D) 100% PC, and (E) without lipids. (F) Deconvolution of the spectra at t = 0.

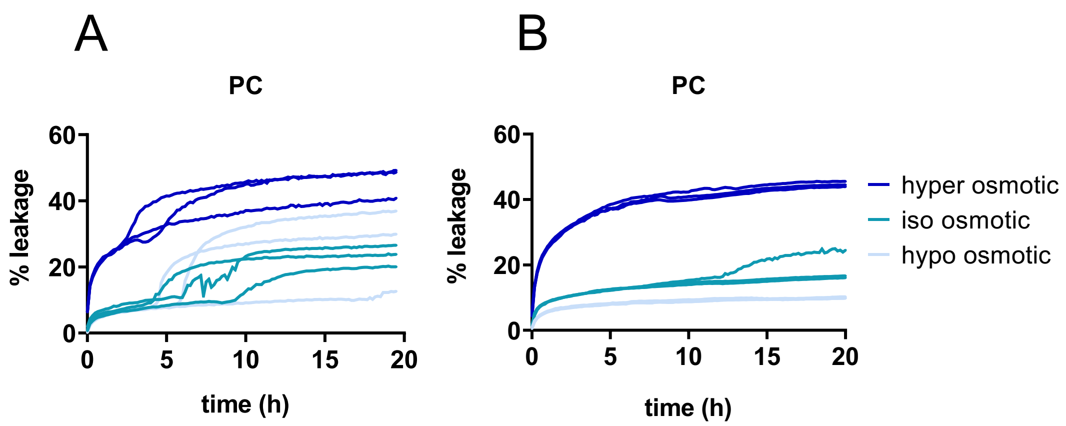


**Figure S4.** Additional datasets of hIAPP induced leakage of pure PC membranes. (A, B) Independent experiments demonstrating the variability in the onset of aggregation-induced leakage. In (B), most of the individual traces do not reach the second phase of leakage during the time course of the experiment.

**Table S1.** Hydrodynamic radii of the different LUV preparations as found by dynamic light scattering.

| Fluorophore | Lipid composition | Hyperosmotic | Isoosmotic | Hypoosmotic |
|---|---|---|---|---|
| Calcein | PC / PS | 185.8 ± 2.0 nm | 185.3 ± 0.8 nm | 185.2 ± 1.7 nm |
| | PC / PG | 182.1 ± 1.4 nm | 182.6 ± 2.9 nm | 181.1 ± 2.5 nm |
| | PC / CL | 192.3 ± 0.6 nm | 187.7 ± 1.8 nm | 191.5 ± 1.3 nm |
| | PC | 181.7 ± 4.7 nm | 182.5 ± 4.6 nm | 176.8 ± 2.6 nm |
| Carboxyfluorescein | PC / PS | 233.7 ± 1.5 nm | 228.7 ± 3.3 nm | 204.6 ± 2.0 nm |
| | PC / PG | 195.0 ± 2.6 nm | 195.2 ± 0.8 nm | 185.3 ± 1.1 nm |
| | PC / CL | 236.0 ± 2.2 nm | 226.7 ± 3.5 nm | 214.0 ± 0.6 nm |
| | PC | 188.0 ± 2.2 nm | 184.5 ± 1.0 nm | 176.1 ± 0.7 nm |
| None | PC / PS | n.a. | 149.8 ± 2.9 nm | n.a. |
| | PC / PG | | 148.5 ± 2.8 nm | |
| | PC / CL | | 154.7 ± 3.0 nm | |
| | PC | | 175.9 ± 11.6 nm | |
